# Verification of emerging genomic mutations in *Mycobacterium tuberculosis* allows transmission chains to be distinguished in an epidemiological typing cluster extending over thirty years

**DOI:** 10.1101/2025.02.07.637016

**Authors:** Rina de Zwaan, Gerard de Vries, Ella Ubbelohde, Arnout Mulder, Miranda Kamst, Karin Rebel, Saskia Kautz, Kristin Kremer, Richard M Anthony, Dick van Soolingen

## Abstract

Whole genome sequencing (WGS) is able to identify epidemiological links between *Mycobacterium tuberculosis* isolates. Recent clustering can be ruled out using a pre-defined single nucleotide polymorphism (SNP) threshold. If WGS clusters grow significantly over time limited genetic variability hampers epidemiological investigations.

Newly emerging (informative) SNPs in isolates of an extended cluster growing for more than 30 years to >150 cases in the Netherlands were analysed. WGS data was analyzed from 61 sequencing files from 54 patients. Genomic positions that varied within the cluster isolates were carefully screened for minority populations in other isolates from the cluster. A transmission scheme was generated on the basis of WGS data alone then compared to the epidemiological information available.

Fifty-two informative SNPs were identified, eight of which were also detected as mixed variants. One emerging SNP in *dnaA* (1199G>A R400H) has been observed in other transmitted strains and may be under selection. There was high concordance between the transmission chains suggested on basis of the newly emerging SNPs and scenario’s identified using classical epidemiological cluster investigations.

Analysis of filtered SNPs accumulating in the genome of *M. tuberculosis* in large clusters contains information on transmission dynamics and can be used to support epidemiological investigations.

## Introduction

In the Netherlands, all *Mycobacterium tuberculosis* complex isolates have been subjected to epidemiological typing by whole genome sequencing (WGS) at the national tuberculosis reference laboratory since 2016 [1]. Isolates within a 12 single nucleotide polymorphism (SNP) distance are reported as ‘clustered’ to the Municipal Public Health Services (MPHs) who perform interviews to identify and confirm epidemiological links [2]. However, most international WGS analysis pipeline’s nowadays apply a stricter threshold of six SNPs to rule in a possible recent transmission [3–4]. Furthermore, SNP filtering in our pipeline is less strict than in most other pipelines, which results in an increased sensitivity to genetic variation and a parallel reduction in specificity (overcalling of SNPs). We estimate the increased SNP calling in our pipeline to be between 0 and 3 SNPs based on observations in international quality control exercises [1, 5].

In low transmission settings such as the Netherlands epidemiological clusters mostly consist of two to three cases, but occasionally large, longer-term clusters occur. In general, the detectable genetic variability between *M. tuberculosis* isolates in even large transmission clusters is limited [6–8]. Moreover, SNP variability between isolates from a single individual can be as large as between isolates from different individuals in a cluster [5, 9] Nonetheless, as the population structure of *M. tuberculosis* is clonal, in principle a higher resolution can be achieved by tracking truly emerging SNPs, as non-fixed alleles in a part of the isolates. This approach should enable a more precise reconstruction of transmission chains [10–13]. This was already suspected in 2010 by investigating single colony cultures of *M. tuberculosis* isolates associated with the large Harlingen outbreak in the Netherlands; the occurrence of new mutations in a part of the colonies of a sub set of the isolates was proven by PCR and could be used to distinguish different sources of spread in the extending cluster [10]. Thus if SNP calling is sufficiently accurate, a higher resolution in typing can be achieved by following the accumulation of specific emerging SNPs and tracking them as fixed or non-fixed alleles in isolates within transmission chains.

Here, we investigate WGS data of the second largest typing cluster in the Netherlands that has been followed by three different molecular typing methods (initially IS*6110* RFLP, then VNTR-MIRU 24, then WGS) over a time period of 30 years [2], with the objective to study the accumulation of new SNPs and to test their utility to distinguish transmission forks introduced by subsequent sources of spread. The predicted evolutionary branching of the strain was compared to information available at MHSs on possible sources of spread in the cluster and subsequent cases, in order to investigate the utility of the cluster sub-branching information in the epidemiological investigation of MHSs.

## Methods

Since 1993, DNA fingerprinting of all *M. tuberculosis* isolates has been performed in the Netherlands, using RFLP typing from 1993-2008, VNTR-MIRU from 2004-2018 and WGS from 2016 to the present. TB public health nurses systematically investigate possible epidemiological links between clustered cases. This activity, the so called ‘cluster investigation’, is prioritised for recent transmission defined as cases that occur within two years after another case. A link is confirmed if the investigation reveals that the newly clustered case was indeed in contact with another clustered and infectious case (e.g. a family member, friend) or if the person was at a particular location at the same time with a clustered infectious case. Since 2009, results of cluster investigations are reported to the National TB Register (NTR), including the identified source case.

A single large cluster, that existed throughout the whole epidemiological typing period since 1993 and with a substantial number of confirmed epidemiological links between clustered cases, was selected for detailed study. We requested the NTR Registration Committee to provide demographic data of cases in this cluster and the results of cluster investigations. All isolates in this cluster had been typed with WGS since 2016. Additionally, patients from before 2016 with a confirmed epidemiological link were selected for re-typing by WGS. MPHs provided information from routine cluster investigation on the most likely place of transmission of patients in the cluster with a confirmed epidemiological link. The laboratory technicians were blinded to information on possible epidemiological links between the selected isolates.

From the cluster, 61 WGS sequencing files of *M. tuberculosis* isolates from 54 individuals were analysed. From four individuals the sequences from two different isolates were included; two individuals had two isolates (<6 months apart) and two others had two isolates as a result of a reactivation (>6 months apart). An additional three isolates were included twice as duplicate controls for the WGS sequencing and data analysis procedure.

The isolates were sequenced using an Illumina (CA, USA) HiSeq^TM^ 2500, NextSeq^TM^ 500 or NextSeq^TM^ 550 sequencer. Unpaired reads were mapped against the H37Rv reference genome using Bowtie2, (GenBank accession: AL123456.3). Breseq v0.28.1 was used to detect SNPs and INDELs (insertions and deletions). The in-house pipeline called SNPs using a ≥80% allele frequency cut off [1]. Called SNPs were excluded from the SNP distance calculation when present in ‘difficult to sequence’ genomic regions according to an exclusion list or when they were within 12 bp from another SNP [3]. All sequencing data were subjected to our routine quality check, specifically an average coverage of 50x or more, no contamination detected (i.e. <3 SNPs called in ribosomal genes *rrs* or *rrl* genes and no more than five specific reads of another lineage in any of the 62 Coll SNPs [14]). Also 15 high confidence first line resistance mutations (supplementary Table S2) were screened for mutant reads at <80% to exclude a mixed resistance genotype / emerging resistance. The sequencing data generated is available online, NCBI accession numbers are listed in supplementary table S1.

### Additional Analysis & WGS transmission scheme

Any SNPs not present in all sequenced isolates from the cluster called by our pipeline vs the reference genome were graded ‘confident’ or ‘non-confident’ based on visual inspection of the SAMtools pile ups [15]. These SNPs were then screened for their distribution in the Dutch WGS database (containing at the moment of screening 3,981 isolates from 53 different lineages [14]). Allele frequencies for any confident SNPs not present in all sequenced isolates in the cluster were then examined in all the isolates from the cluster sequenced. These allele frequencies were used to identify emerging SNPs; an allele frequency over 95% was considered ‘fixed’ and below 5% as wild type, an allele frequency between 5% and 95% was considered a mixed SNP.

A WGS transmission scheme was then produced based on the distribution of all ‘confident’ (both fixed and mixed) SNPs based on the assumptions that;

i. SNP acquisition is unique, i.e. the same SNP did not occur randomly twice.
ii. Fixed SNPs are not lost in the transmission scheme.
iii. Mixed SNPs are either transmitted or not transmitted to a secondary case.

The transmission scheme, generated from the WGS data was then compared to the transmission inferences obtained on the basis of cluster investigations and the known relationships between the isolates (including duplicates and multiple isolates from the same individual).

### Ethical clearance

The NTR Registration Committee approved our request and provided the WGS and epidemiological data. As the routine collected data were pseudonymized, the study was not subjected to the Dutch law for Medical Research Involving Human Subject Act (WMO).

## Results

### Cluster description based on epidemiological information

The selected cluster was identified in 1993 and composed of 157 tuberculosis cases by the end of 2022 (overview in figure S1). In February 2003, infectious TB was diagnosed in a new patient (case 5, Figure 1,transmission chain 1; red) who frequently visited pubs in one specific street in Rotterdam. This patient was lost to follow-up during treatment and traced in November 2003, being again highly infectious (case 23). Between 2003-2014, the cluster increased by 121 patients; 80 (66%) were living in the city of Rotterdam, of which 30 (38%) reported frequent visits to the pubs in the mentioned street. A new transmission chain occurred in Amsterdam (Figure 1 transmission chain 7; dark blue Rotterdam, light blue Amsterdam) with two highly infectious cases. Case 96 was diagnosed in 2009 by contact investigation around a child with TB (case 94). A contact of this person developed tuberculosis in 2012 (case 114) and had recurrent tuberculosis (case 129), more than two years after completing the first treatment. Several new transmission locations were identified by cluster investigation, such as a pub in another town (3 cases; not shown), a penitentiary institute (3 cases), a coffee shop (5 cases), a school (4 cases, Figure 1, transmission chain 6; purple) and a hospital (one case of nosocomial transmission). Case 153 developed tuberculosis nine years after being diagnosed with a TB infection in a contact investigation and having received preventive treatment (Figure 1, transmission chain 5; green).

**Figure 1.**
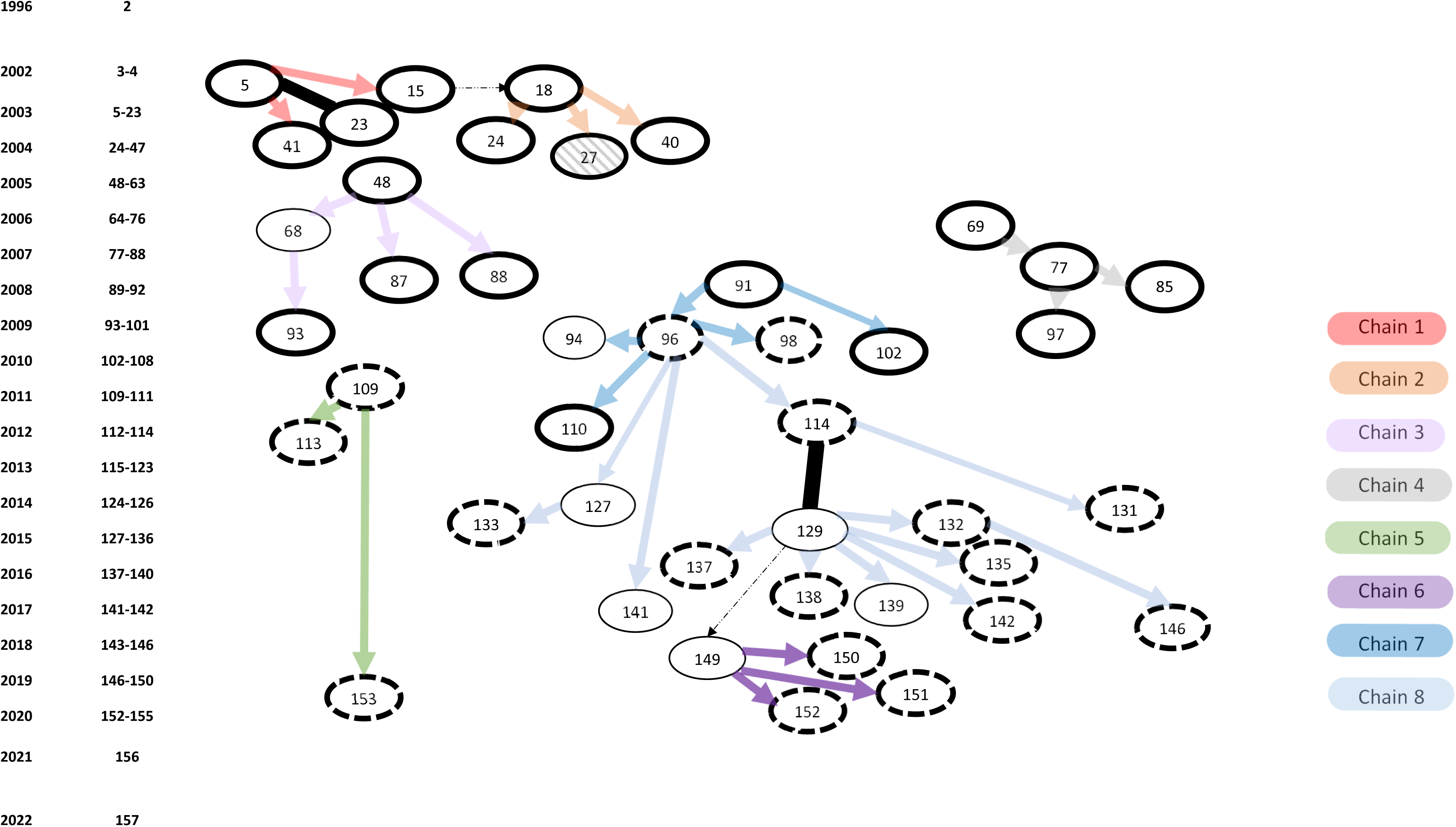
Transmission scheme based on epidemiological information. Cases sequenced with an epidemiological link confirmed by MHSs are present as ovals, with arrows showing the direction of transmission. Relapse cases in the same individual are linked with a solid black line. Oval borders indicate the location of the case; solid thick border Rotterdam; dashed thick border Amsterdam; thin border; other towns or villages in the Netherlands. The arrow colour indicates a distinct epidemiological transmission chain. Dotted arrows indicate situations in which transmission was considered likely. The cases and epidemiological transmission chains are presented using the same colours in Figure 2 (except case 27 shaded, for which WGS data was unanalysable due to contamination of the culture). The year and number of isolates from the cluster is indicated on the left hand side (and in table S1).

### Cluster description based on WGS data (SNPs)

A total of 773 SNPs vs the reference genome was called by our pipeline in all the sequenced isolates from the selected cluster, 706 of these SNPs were shared by all 61 WGS files (available online Table S1) investigated from the cluster on basis of WGS and uninformative. All duplicate controls showed exactly the same confident SNPs as fixed, wild type or mixed. Of the remaining 67 SNPs 53 were manually graded as confident, 14 as non-confident (Table 1). One SNP (Table 1, SNP 64), which was confidently graded, was present in all members of the cluster but not always reliably called by our pipeline; this SNP was thus considered uninformative and not used in the construction of the transmission scheme. The final analysis was thus based on 52 SNPs (Table 1).

**Table 1.**
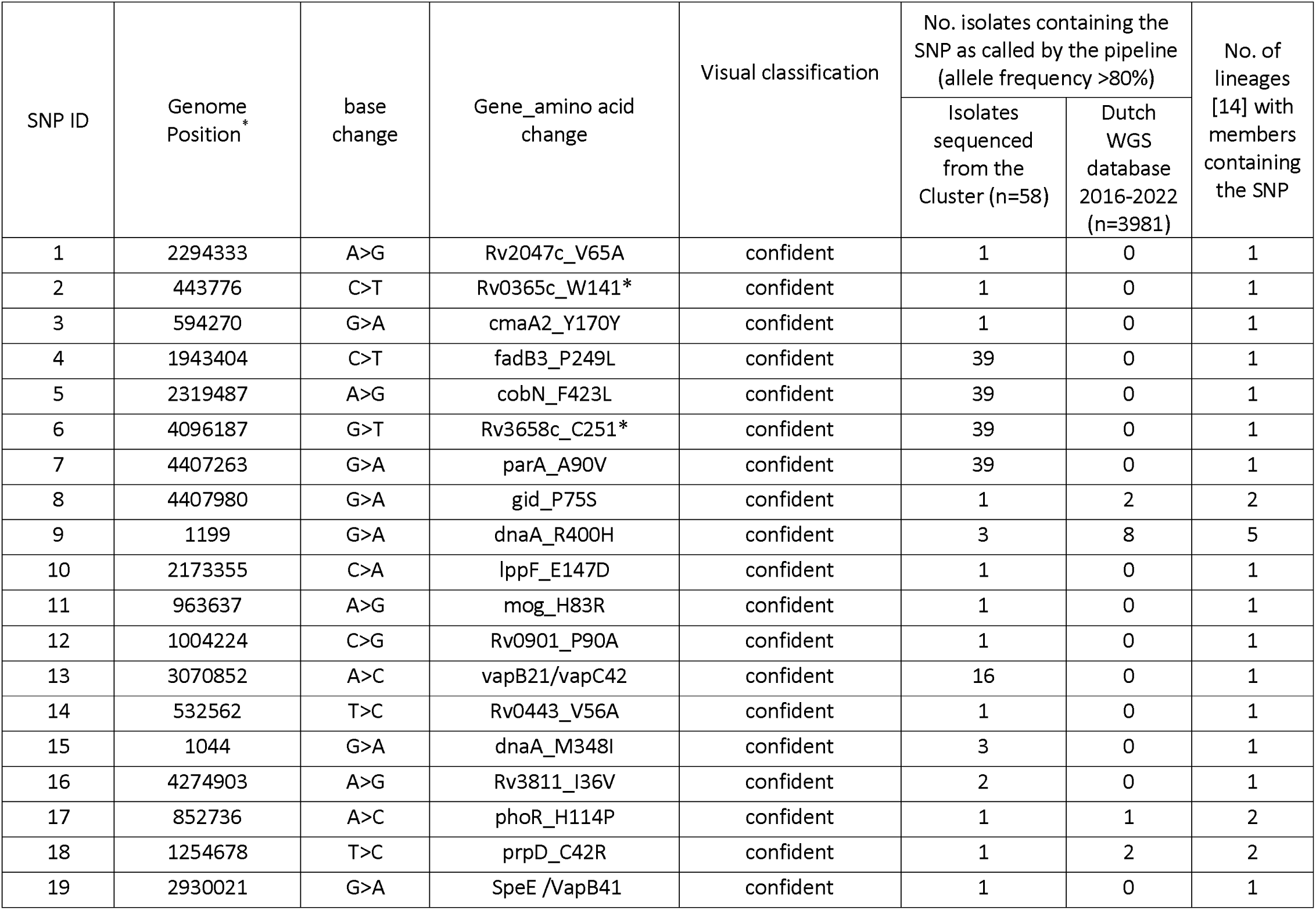
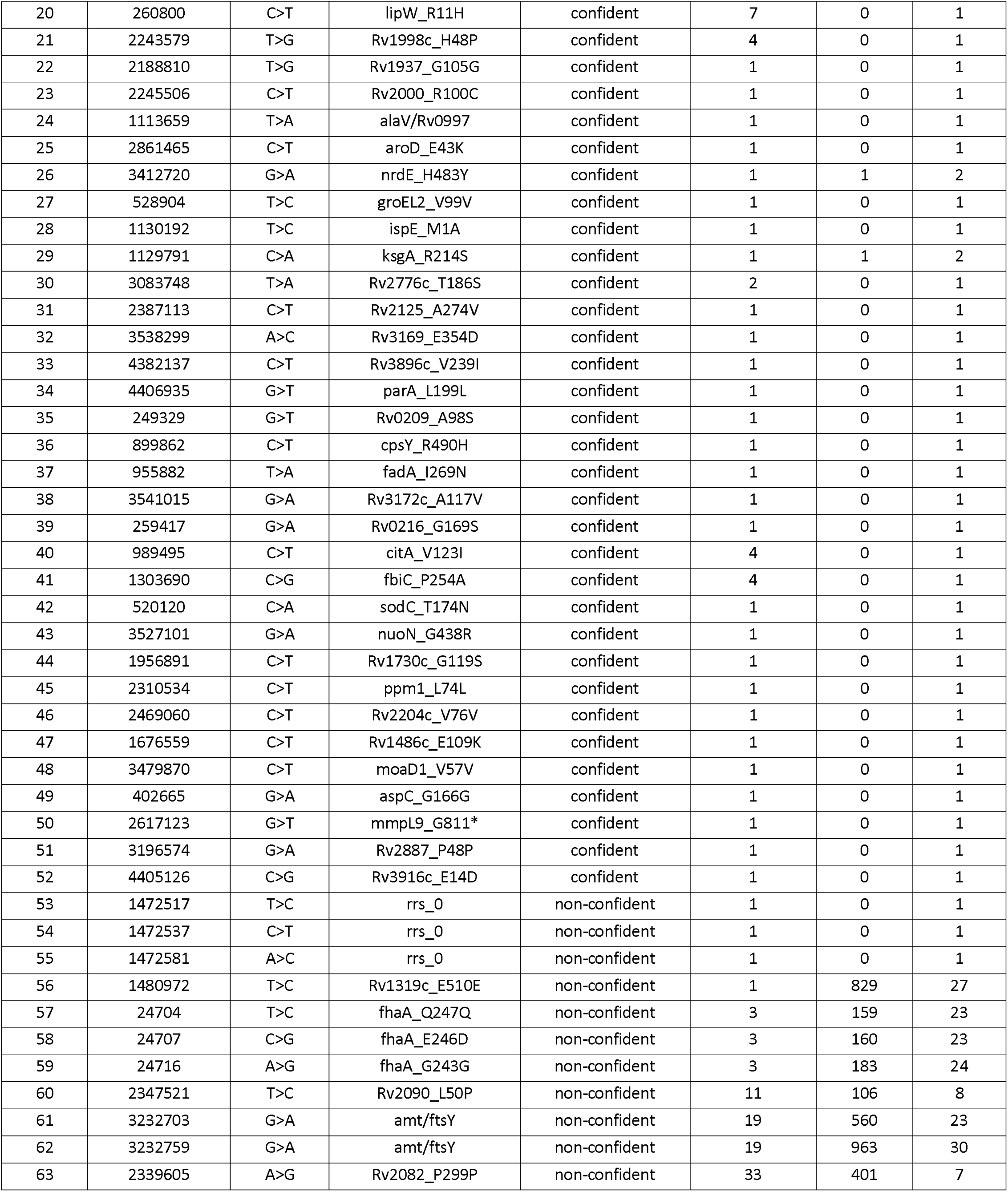
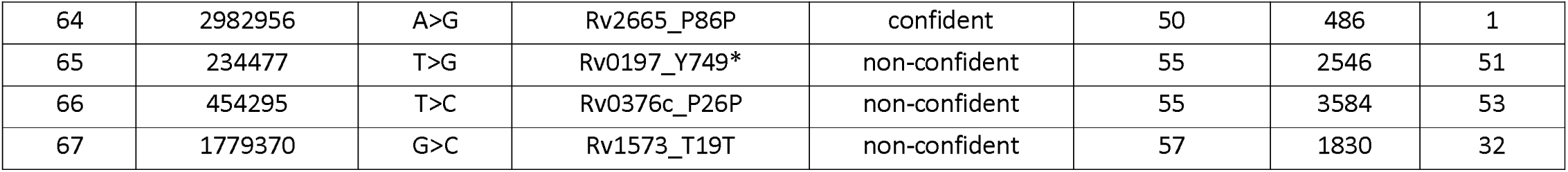
Overview of all SNP diversity detected in the cluster, the visual grading of the SNPs and the distribution of the diverse identified in the clustered isolates in the NL database of 3981 isolates. . * = relative to the reference genome H37Rv (AL123456.3)

| SNP ID | Genome Position * | base change | Gene_amino acid change | Visual classification | No. isolates containing the SNP as called by the pipeline (allele frequency >80%) |  | No. of lineages [14] with members containing the SNP |
| --- | --- | --- | --- | --- | --- | --- | --- |
|  |  |  |  |  | Isolates sequenced from the Cluster (n=58) | Dutch WGS database 2016-2022 (n=3981) |  |
| 1 | 2294333 | A>G | Rv2047c_V65A | confident | 1 | 0 | 1 |
| 2 | 443776 | C>T | Rv0365c_W141* | confident | 1 | 0 | 1 |
| 3 | 594270 | G>A | cmaA2_Y170Y | confident | 1 | 0 | 1 |
| 4 | 1943404 | C>T | fadB3_P249L | confident | 39 | 0 | 1 |
| 5 | 2319487 | A>G | cobN_F423L | confident | 39 | 0 | 1 |
| 6 | 4096187 | G>T | Rv3658c_C251* | confident | 39 | 0 | 1 |
| 7 | 4407263 | G>A | parA_A90V | confident | 39 | 0 | 1 |
| 8 | 4407980 | G>A | gid_P75S | confident | 1 | 2 | 2 |
| 9 | 1199 | G>A | dnaA_R400H | confident | 3 | 8 | 5 |
| 10 | 2173355 | C>A | lppF_E147D | confident | 1 | 0 | 1 |
| 11 | 963637 | A>G | mog_H83R | confident | 1 | 0 | 1 |
| 12 | 1004224 | C>G | Rv0901_P90A | confident | 1 | 0 | 1 |
| 13 | 3070852 | A>C | vapB21/vapC42 | confident | 16 | 0 | 1 |
| 14 | 532562 | T>C | Rv0443_V56A | confident | 1 | 0 | 1 |
| 15 | 1044 | G>A | dnaA_M348I | confident | 3 | 0 | 1 |
| 16 | 4274903 | A>G | Rv3811_I36V | confident | 2 | 0 | 1 |
| 17 | 852736 | A>C | phoR_H114P | confident | 1 | 1 | 2 |
| 18 | 1254678 | T>C | prpD_C42R | confident | 1 | 2 | 2 |
| 19 | 2930021 | G>A | SpeE /VapB41 | confident | 1 | 0 | 1 |
| 20 | 260800 | C>T | lipW_R11H | confident | 7 | 0 | 1 |
| 21 | 2243579 | T>G | Rv1998c_H48P | confident | 4 | 0 | 1 |
| 22 | 2188810 | T>G | Rv1937_G105G | confident | 1 | 0 | 1 |
| 23 | 2245506 | C>T | Rv2000_R100C | confident | 1 | 0 | 1 |
| 24 | 1113659 | T>A | alaV/Rv0997 | confident | 1 | 0 | 1 |
| 25 | 2861465 | C>T | aroD_E43K | confident | 1 | 0 | 1 |
| 26 | 3412720 | G>A | nrdE_H483Y | confident | 1 | 1 | 2 |
| 27 | 528904 | T>C | groEL2_V99V | confident | 1 | 0 | 1 |
| 28 | 1130192 | T>C | ispE_M1A | confident | 1 | 0 | 1 |
| 29 | 1129791 | C>A | ksgA_R214S | confident | 1 | 1 | 2 |
| 30 | 3083748 | T>A | Rv2776c_T186S | confident | 2 | 0 | 1 |
| 31 | 2387113 | C>T | Rv2125_A274V | confident | 1 | 0 | 1 |
| 32 | 3538299 | A>C | Rv3169_E354D | confident | 1 | 0 | 1 |
| 33 | 4382137 | C>T | Rv3896c_V239I | confident | 1 | 0 | 1 |
| 34 | 4406935 | G>T | parA_L199L | confident | 1 | 0 | 1 |
| 35 | 249329 | G>T | Rv0209_A98S | confident | 1 | 0 | 1 |
| 36 | 899862 | C>T | cpsY_R490H | confident | 1 | 0 | 1 |
| 37 | 955882 | T>A | fadA_I269N | confident | 1 | 0 | 1 |
| 38 | 3541015 | G>A | Rv3172c_A117V | confident | 1 | 0 | 1 |
| 39 | 259417 | G>A | Rv0216_G169S | confident | 1 | 0 | 1 |
| 40 | 989495 | C>T | citA_V123I | confident | 4 | 0 | 1 |
| 41 | 1303690 | C>G | fbiC_P254A | confident | 4 | 0 | 1 |
| 42 | 520120 | C>A | sodC_T174N | confident | 1 | 0 | 1 |
| 43 | 3527101 | G>A | nuoN_G438R | confident | 1 | 0 | 1 |
| 44 | 1956891 | C>T | Rv1730c_G119S | confident | 1 | 0 | 1 |
| 45 | 2310534 | C>T | ppm1_L74L | confident | 1 | 0 | 1 |
| 46 | 2469060 | C>T | Rv2204c_V76V | confident | 1 | 0 | 1 |
| 47 | 1676559 | C>T | Rv1486c_E109K | confident | 1 | 0 | 1 |
| 48 | 3479870 | C>T | moaD1_V57V | confident | 1 | 0 | 1 |
| 49 | 402665 | G>A | aspC_G166G | confident | 1 | 0 | 1 |
| 50 | 2617123 | G>T | mmpL9_G811* | confident | 1 | 0 | 1 |
| 51 | 3196574 | G>A | Rv2887_P48P | confident | 1 | 0 | 1 |
| 52 | 4405126 | C>G | Rv3916c_E14D | confident | 1 | 0 | 1 |
| 53 | 1472517 | T>C | rrs_0 | non-confident | 1 | 0 | 1 |
| 54 | 1472537 | C>T | rrs_0 | non-confident | 1 | 0 | 1 |
| 55 | 1472581 | A>C | rrs_0 | non-confident | 1 | 0 | 1 |
| 56 | 1480972 | T>C | Rv1319c_E510E | non-confident | 1 | 829 | 27 |
| 57 | 24704 | T>C | fhaA_Q247Q | non-confident | 3 | 159 | 23 |
| 58 | 24707 | C>G | fhaA_E246D | non-confident | 3 | 160 | 23 |
| 59 | 24716 | A>G | fhaA_G243G | non-confident | 3 | 183 | 24 |
| 60 | 2347521 | T>C | Rv2090_L50P | non-confident | 11 | 106 | 8 |
| 61 | 3232703 | G>A | amt/ftsY | non-confident | 19 | 560 | 23 |
| 62 | 3232759 | G>A | amt/ftsY | non-confident | 19 | 963 | 30 |
| 63 | 2339605 | A>G | Rv2082_P299P | non-confident | 33 | 401 | 7 |
| 64 | 2982956 | A>G | Rv2665_P86P | confident | 50 | 486 | 1 |
| 65 | 234477 | T>G | Rv0197_Y749* | non-confident | 55 | 2546 | 51 |
| 66 | 454295 | T>C | Rv0376c_P26P | non-confident | 55 | 3584 | 53 |
| 67 | 1779370 | G>C | Rv1573_T19T | non-confident | 57 | 1830 | 32 |

Subsequent comparison of the distribution of all 67 variable SNPs in the entire Dutch WGS database showed that all but three of the 14 SNPs manually graded as non-confident SNPs were randomly distributed throughout the database, and present in more than 100 unrelated isolates spread over at least seven different lineages, as a result of poor mapping/sequencing at these positions (Table 1). The non-confident SNPs e.g. included three SNPs in the *fhaA gene* (Table 1, SNPs 57-59) which had already been removed from our routine cluster analyses by the pipeline based on the fact they were within 12 base pairs from each other.

In total 46 of the 52 SNPs manually classified as ‘confident’ were exclusive to the cluster in the Dutch WGS database. Five SNPs (Table 1; SNPs 8, 17, 18, 26, 29) were only shared by one or two isolates from one different lineage in the Dutch WGS database. One confidently graded SNP in *dna*A (Table 1, SNP 9) was shared by eight other isolates from four different lineages.

The accumulation of the 52 confident SNPs in a proportion of the isolates was used to inform the WGS-based transmission scheme (Figure 2) based on the assumptions outlined in the methods.

**Figure 2:**
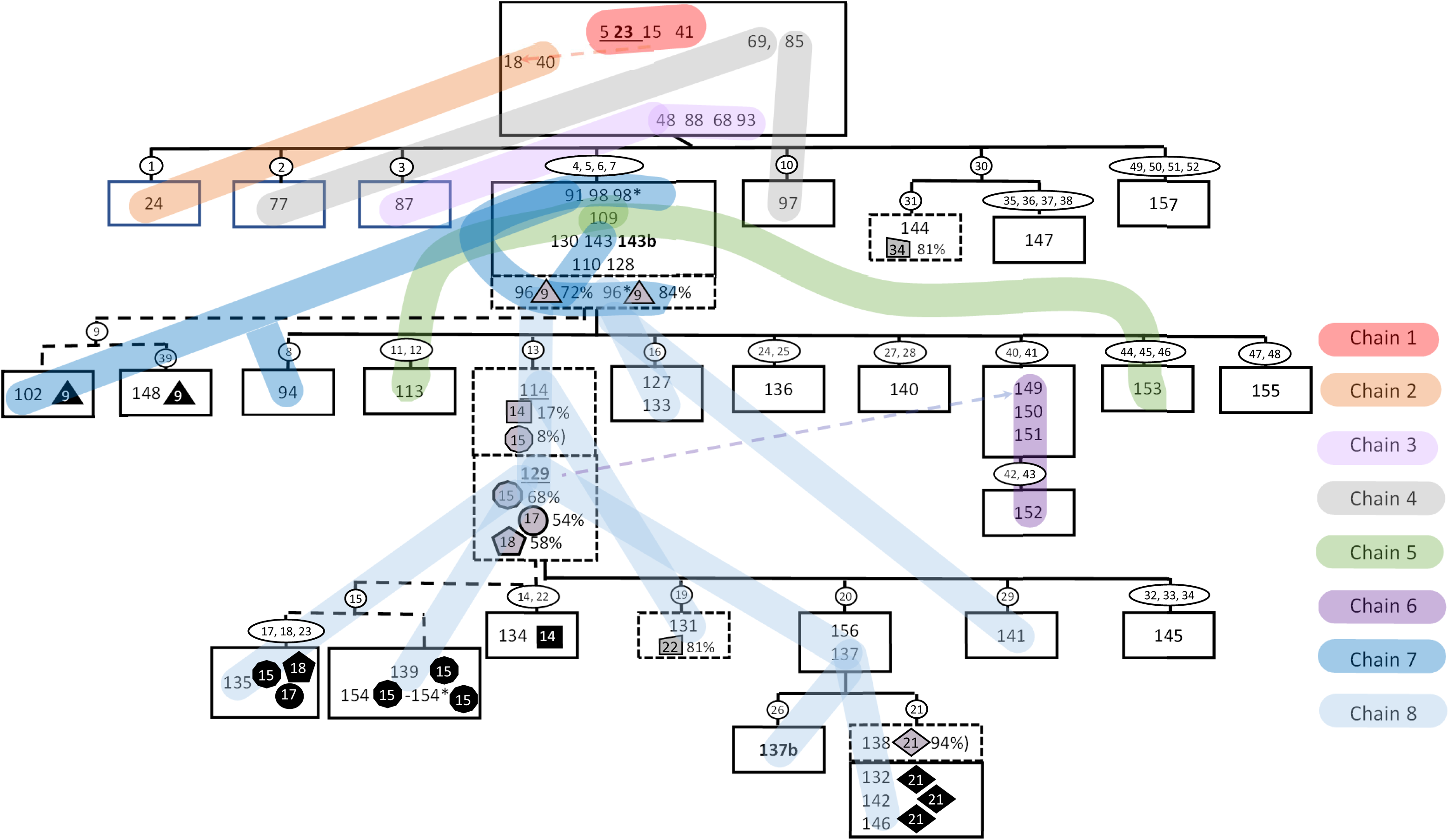
Transmission scheme based on WGS data. Numbers inside boxes indicate NTR cases (Figure 1), cases with zero fixed SNP differences are inside the same box. NTR case numbers appearing more than once indicate duplicate sequences generated from the same isolate are indicated by an *. Follow up isolates from the same NTR case are indicated in bold with a “b”. Reactivations have a different NTR case number but are underlined with the second episode in bold. Numbers inside white ovals indicate a fixed SNP (Table 1); unfixed SNPs inside shaded symbols; a grey symbol if the SNP was mixed and a black symbol when in subsequent cases where the SNP was fixed, they have their frequency indicated as a percentage after a comma inside the box. Solid lines linking boxes indicate the inferred transmission of fixed SNPs, dashed lines the inferred transmission of mixed SNPs. Coloured lines indicate isolates in epidemiologically linked transmission chains(Figure 1).

There were 44 confident SNPs that were not present in all isolates in the cluster and were fixed (>=95% of reads) in all the files in which they were observed. The remaining eight confident SNPs were present in at least one file as a mixed SNP (<95% of the reads). Six of the eight mixed SNPs were fixed in at least one isolate (Figure 2, grey [mixed] and black [fixed] symbols). The remaining two SNPs were only found as mixed in only a single isolate and never fixed (SNPs 22 and 34).

The duplicate controls were identical and placed together on the WGS transmission tree with pair 96 both containing mixed SNP 9 (Figure 2, triangle symbol) at 72% and 84% respectively.

Only two of the eight mixed SNPs were present as a mixed SNP in more than one sequencing file; one duplicate control (SNP 9 in sample 96 mentioned above) and one (SNP 15, Figure 2 round symbols) in two different sequencing files. The two files with a shared mixed SNP 15 were serial isolates from the same individual (with a reactivation) (Figures 1 and 2; 114, 129). These results were derived from isolates collected in September 2012 (SNP 15; 8%) and March 2015 (SNP 15; 68%) from the same patient. Four files also contained a fixed SNP 15(135, 139, 154 and 154*); two of these were duplicate controls (154, 154*). The first isolate in which SNP 15 was mixed (September 2012) also contained a mixed SNP 14 (17%). SNP 14 was fixed in one additional isolate (134) without SNP 15. In the second isolate (Figure 2 129; March 2015) two other additional mixed SNPs were present (SNP 17; 54%, SNP 18; 58%). One isolate (Figure 2; 135) contained both SNPs 17 and 18 fixed along with SNP 15.

### Comparison of the epidemiological and WGS based transmission schemes

The comparison of the information from the WGS analysis and the epidemiological investigations included 33 confirmed epi-links (68 cases) and two relapses. The serial isolates case 5 and its relapse (case 23) were identical, while the isolates from case 114 and its relapse (case 129) contained mixed SNPs; (SNP 14; 17%, and SNP 15; 8%, in the first episode SNP 15; 68%, SNP 17; 54% and SNP 18; 58% in the second isolate (Figure 2).

WGS failed for one isolate selected for WGS analysis (Figure 1, case 27 shaded oval). Out of the remaining 33 of the confirmed epi-links and two repeat isolates screened for SNPs, 16 were identical, 11 contacts gained one SNP, four contacts gained two SNPs and two contacts gained three new confident SNPs (Table 2). There were two epi-linked isolates (cases 85 and 97) that based on the WGS scheme lost a unique fixed SNP identified in the isolate of putative source case 77 (Table 2, Figures 1 and 2). In retrospect it was concluded that these two cases (85, 97) were related cases to case 77 but could also very well have been infected by case 69, who was the source of infection of case 77, which would be consistent with the WGS transmission scheme (Figure 2; supplementary table S1).

**Table 2.** Distribution of SNP distances between isolates considered to have an epidemiological link based on contact investigations undertaken by the MHSs.

| SNPs between source and secondary case |  | Number of Epi-links |
| --- | --- | --- |
| SNPs gained | SNPs Lost |  |
| 0 | 0 | 16 |
| 1 | 0 | 10 |
| 2 | 0 | 5 |
| 3 | 0 | 2 |
| 0 | 1 | 1 |
| 1 | 1 | 1 |

All cases that occurred since 2015 in patients residing in Amsterdam and surrounding areas contained SNPs 4-7 and for most epidemiological links were identified (Figure 1 and 2, transmission chain 7 dark and light blue lines).

## Discussion

This study shows that with proper verification of variable mutations identified in a large cluster useful information was revealed with respect to the micro-evolution of the strain. The resulting branching divided the large cluster in several satellite-clusters, which is relevant for monitoring the development of the outbreak and to steer public health actions. The inferences based on a detailed WGS analysis were almost entirely compatible with the possible scenarios developed by the MHSs based on information extracted from epidemiological investigations. The initial outbreak occurred in Rotterdam, the second largest city in the Netherlands, and was associated with known risk factors like frequent visits to pubs in one area. From 2009 onwards, cases emerged in a sub-cluster in Amsterdam. Over time the outbreak strain acquired four confident SNPs, which were present in isolates linked to subsequent cases in Amsterdam, but not in the initial cases in Rotterdam.

We classified all 67 SNPs that were variably present in the cluster using two independent approaches; firstly visual inspection of the sequencing reads (SAMtools Pileups [15]) and secondly an analysis of distribution of the SNPs in the WGS database in the Netherlands comprising 3,981 diagnostic isolates (Table 1). Of the 14 visually non-confident called SNPs, 11 were also identified as non-informative based on their presence in a large number of genetically diverse isolates. A proportion of these SNPs were found in genomic regions that have been previously recognized as poorly mapping [16]. The remaining three SNPs in *rrs* (Table 1; SNP 53-55) were considered unreliable, presumably as a result of contamination with other bacteria. Finally, one confidently called SNP (64; Rv2662_P86P) was found in 486 other isolates in the WGS database, all belonging to one lineage and was likely present in all members of this lineage but not called by our pipeline in a small proportion of lineage 4.1.2.1 isolates due to poor mapping of a A>G substitution directly next to a string of G nucleotides (Table 1).

Six of the 52 remaining confident SNPs were not unique to the cluster but present in one, two and in one case eight additional isolates in the WGS database of 3,981 unique isolates. Positions under selection in clonal organism such as *M. tuberculosis* are subject to homoplasy, most notably mutations associated with drug resistance [17]. These six SNPs are thus potential candidates for positions under selection. Strikingly, SNP 9 [dnaA_R400H] emerged in the cluster and was identified in eight unrelated isolates in diverse lineages, present in the WGS database (Table 1). Additionally, a second SNP also emerged in this cluster in the *dna*A gene (M348I), this SNP was only identified in isolates from the cluster studied (Table 1, SNP 15). Mutations in *dna*A have previously been observed in spontaneous *M. tuberculosis* drug resistant laboratory mutants [11], and have been associated with increased survival during INH treatment and more robust growth at the MIC of isoniazid [18]. Furthermore, *dna*A mutations have also been observed in more efficiently transmitted variants in another cluster of isolates that was subject to detailed SNP analysis [12]. Notably, both *dna*A mutations observed in this cluster were initially identified as mixed SNPs and transmitted to other individuals. Furthermore, the *dna*A_M348I mutation was also present in two isolates from a treatment failure / reactivation case (cases 114, 129) isolated more than two years apart (Figure 2, transmission chain 7; blue) with at least four mixed SNPs which were identified as fixed in other individuals who were presumably infected by this case.

After filtering, either manually or screening our database for unexpected homoplasy, high confidence informative SNPs duplicate controls were identical and serial/reactivation isolates were all directly linked in the WGS scheme.

We identified the SNP variability within the cluster on the basis of SNP thresholds and then filtered this short list of SNPs based on a screen for variability of these variable positions in a larger database allowing “noisy” SNPs to be identified and filtered out of the analysis. In this way true potentially epidemiologically informative SNPs within the cluster were identified. The resulting short list of confident informative positions could then be analysed in all other isolates sequenced from the cluster in detail with the aim of identifying the isolates in the cluster in which the SNPs were emerging/not fixed [10–12, 19]. This approach in cancelling sequence noise using a large database to screen for epidemiologically informative variation, adds to the epidemiological typing. Comparing the epidemiological data available to WGS based transmission scheme supports the assumption that careful analysis can identify epidemiologically informative variability in closely related sequences.

### Limitations

Even with a permissive SNP calling algorithm for a significant proportion of individuals (20 of 54 ,37%) in the cluster analysis we were unable to identify any confident SNP variation with at least one other case. Long read sequencing or the use of pan genome analysis may allow the analysis of more variable regions of the genome that currently cannot be confidently analysed using short read sequencing [16] allowing the detection of large genome rearrangements as well as potentially allowing analysis of regions not present in the reference genome currently used could potentially increase the typing resolution [20, 21] within recently clustered isolates. We sequenced cultured isolates raising the possibility that sub-populations of bacteria present in the sample may grow at different rates, be lost, or additional mutations accumulate during culture [22]. Finally, we did not screen the whole genomes for mixed SNPs to streamline the analysis, targeted only SNPs that were variable/informative within the clustered isolates sequenced. This detailed analysis was performed on a single larger cluster. The identification of emerging SNPs will only be informative when the majority of cases in the cluster are identified and sequence data is available for at least the majority of the potentially linked cases.

## Conclusion

In conclusion, although clustering can be effectively ruled out on the basis of SNP distance thresholds alone we demonstrate there is additional information available in micro variability within the WGS data from isolates in a large cluster that can be useful to identify possible links and target epidemiological investigations. We confirm mixed SNPs, if present, may be fixed in subsequent isolates and strongly suggest a transmission direction and direct epidemiological link as supported by the epidemiological information available from the cluster studied and recently proposed by others [19, 23, 24]. Screening for these mixed SNPs in a long established cluster can therefore provide epidemiologically useful information. Strategies to automate and validate this type of SNP analysis could be implemented. Marker SNPs characteristic of specific transmission chains [25] can be identified in larger clusters and are potentially more precise than SNP thresholds to identify epidemiologically linked isolates.

## Acknowledgments

This project was funded by the Dream Fund project “No more pandemics” of the Dutch Postcode Lottery, and the Dutch Ministry of Health, Welfare and Sports. The funders had no role in the study design, data collection and interpretation, or the decision to submit the work for publication. We would also like to acknowledge the support of The Netherlands Organisation of Health, Research and Development (ZonMw project 541002006) which supported the development of the SNP screening algorithm. We thank the MHSs in the Netherlands for their highly valuable contribution to this paper and ongoing collaboration.

## Contributions (CRediT)

Rina de Zwaan: Data curation, Formal Analysis, Writing review and editing.

Gerard de Vries: Conceptualization, Data curation, Methodology, Writing – review & editing.

Ella Ubbelohde: Data analysis, Writing – review & editing.

Arnout Mulder: Data curation.

Miranda Kamst: Project administration, Supervision.

Karin Rebel: Data curation.

Saskia Kautz: Data curation.

Kristin Kremer: Funding acquisition, Project administration, Supervision, Writing – review & editing.

Richard Anthony: Conceptualization, Funding acquisition, Methodology, Supervision, Writing – original draft.

Dick van Soolingen: Conceptualization, Funding acquisition, Writing -original draft, Writing – review & editing.

